# Activity dependent regulation of Angiomotin modulates dendritic complexity in hippocampal neurons

**DOI:** 10.64898/2026.10.26.761988

**Authors:** Michael Wigerius, Stefan Krueger, James P. Fawcett

## Abstract

Neuronal activity alters synaptic protein composition, which is important for maintaining dendritic complexity. Multiple signaling networks have been implicated in these processes, but how increased neuronal activity regulates their expression remains incompletely understood. The long isoform of Angiomotin, AMOT-130, is a scaffolding protein that recruits signaling proteins involved in regulating cell morphology and has been implicated in dendritic spine morphogenesis during early development. Here, we show that RNAi-mediated depletion of AMOT-130 reduces dendritic arbor complexity in neurons as early as DIV9. Elevated neuronal activity similarly decreases dendritic complexity which is accompanied by reduced AMOT-130 levels. Overexpression of AMOT-130 counteracts this morphological effect, suggesting that its activity-dependent downregulation contributes to dendritic remodeling during early development. We further, demonstrate that elevated neuronal activity promotes ubiquitin-mediated proteasomal degradation of AMOT-130, identifying a mechanism regulating its abundance in developing neurons. These findings suggest that maintaining sufficient AMOT-130 protein levels is necessary for normal dendritic arborization.

## Introduction

Most neurons in the brain share extensive dendritic arbors that enable them to make new synaptic connections during the development. During development, genetic programs, axon guidance cues, and cell-adhesion molecules work together to direct neuronal growth, target selection, and the formation of appropriate synaptic connections (Lefebvre, Sanes, & Kay, 2015). Once developed, neuronal circuitry is refined by neuronal activity, leading to synaptic pruning and elimination of dendrites, which are important mechanisms through which neuronal activity shapes developing circuits(Pan & Monje, 2020). These changes in neuronal structure require dynamic regulation of the cytoskeleton through networks of protein-protein interactions that coordinate actin and microtubule organization (Cingolani & Goda, 2008; Dogterom & Koenderink, 2019; Falace et al., 2026; Matus, 2000). Activity-dependent cytoskeletal regulation is also central to synaptic plasticity, in which changes in dendritic spine morphology are necessary for the remodeling of neuronal circuits (Cingolani & Goda, 2008; Nakahata & Yasuda, 2018).

Understanding how activity is translated into changes in morphology is important to define the underlying mechanism driving dendritic and synaptic remodeling. For example, NMDAR activation regulates downstream signaling through calcium mediated events to affect the activity of the small Rho-family of GPTases, Rac, Rho and Cdc42, to control actin dynamics which is critical for both dendritic growth and structural synaptic plasticity (Matsuzaki, Honkura, Ellis-Davies, & Kasai, 2004; Murakoshi, Wang, & Yasuda, 2011; Okamoto, Nagai, Miyawaki, & Hayashi, 2004). In addition, activity has been linked to other intracellular processes including the spatial localization of proteins, protein turnover, and post-translational modifications which all have been shown to regulate dendritic structure and synaptic connectivity (Ehlers, 2003; Matt, Kim, Chowdhury, & Hell, 2019; Vallejo, Codocedo, & Inestrosa, 2017).

One well established mechanism regulating protein abundance is ubiquitination, which involves the attachment of ubiquitin molecules to target proteins to the ubiquitin-proteasome system (UPS) (Bingol & Schuman, 2006). This involves the reversible attachment of the 76-amino acid protein ubiquitin to lysine residues of substrates (Vucic, Dixit, & Wertz, 2011). The attachment of multiple ubiquitin moieties into chains can target proteins for degradation by the multi-subunit proteasome complex, whereas monoubiquitin has been linked to cellular trafficking (Yamada, Yang, & Bonni, 2013).

Among the enzymes that mediate ubiquitin conjugation is the NEDD4 family of E3 ubiquitin ligases, several members of which have been implicated in targeting synaptic proteins in response to activity cues (Ambrozkiewicz & Kawabe, 2015; Widagdo, Guntupalli, Jang, & Anggono, 2017). Conversely, deubiquitinating (DUB) enzymes remove ubiquitin from protein conjugates, counteracting E3 ligase activity and thereby contributing to maintaining overall protein balance (Todi & Paulson, 2011). Such regulation is important for cytoskeletal organization and has been shown to be important for dendritic remodeling (Kuo 2005, 2006). How activity-dependent protein turnover contributes to dendritic architecture remains poorly defined.

Angiomotin (AMOT) is a potential regulator of these processes given it is linked to RhoGTPase regulation and the UPS system in non-neuronal cells (C. Wang et al., 2012; Wells et al., 2006). AMOT encodes two isoforms, AMOT-80 and AMOT-130, which share a C-terminal PDZ-binding motif and a proximal coiled-coil domain. AMOT-130 also contains an N-terminal extension with an F-actin-binding motif and multiple L/PPxY motifs that mediate interactions with Hippo pathway components, positioning this isoform at the intersection of Hippo signaling, cell polarity, and actin dynamics (Y. Wang & Yu, 2024; Wigerius, Quinn, & Fawcett, 2020). We previously demonstrated that AMOT-130 contributes to actin cytoskeletal stability and structural synaptic plasticity in dendritic spines (Wigerius et al., 2018; Wigerius et al., 2020). Furthermore, depletion of both AMOT-130 and AMOT-80 reduces dendritic complexity, but the individual contribution of these isoforms remains unclear (Rojek et al., 2019). AMOT also interacts with proteins that regulate Rho-family GTPase signaling, including GTPase-activating proteins (GAPs) (Wells et al., 2006; Yi et al., 2011), providing a potential link between AMOT, cytoskeletal regulation, and neuronal structure.

Here, we demonstrate that isoform-specific RNAi-mediated depletion of AMOT-130 reduces dendritic arbor complexity, while elevated neuronal activity regulates AMOT-

130 protein levels through ubiquitin-dependent proteasomal degradation involving NEDD4 family ubiquitin ligases. This activity-dependent regulation is functionally linked to dendritic arborization, as the reduction in dendritic complexity induced by elevated activity can be reversed by AMOT-130 overexpression. Taken together, these findings support the idea that global changes in activity control AMOT-130 expression and contribute to homeostatic dendritic remodeling.

## Material and Methods

The YFP-AMOT construct has been described previously (Wigerius et al., 2018). The YFP-AMOT-AAA mutant was generated by site-directed mutagenesis introducing three alanine residues in the N-terminus of AMOT which deletes L/PPxY motif binding (C. Wang et al., 2012). The Myc-NEDD4-1 construct was generated by PCR using HA-NEDD4 (Addgene, #27002) as template and cloned into pcDNA3.1/myc-His B (Thermo Fisher Scientific). The pEGFP-C1 construct was from Taka Bio Inc (Clontech). Isoform specific siRNA oligonucleotides (siRNA#1 and siRNA#2) that target the N-terminal region in AMOT-130 and control (CsiRNA) whose sequence is not homologues to any vertebrate sequence have been described previously (Wigerius et al., 2018).

Primary antibodies used in the study were as follows: goat anti-AMOT-130 that recognizes both AMOT-130 and AMOT-80 isoforms (1:200; C-18, Santa Cruz Biotechnology), mouse anti-myc (1:500, 9E10, Santa Cruz Biotechnology), rabbit anti-2095 (1:1000, (Wigerius et al., 2018), rabbit anti-AMOT-130 for immunostaining (1:1000, ABS1024, EMD Millipore), rabbit anti-NEDD4-1 (1:1000, D82F4, Cell Signaling Technology), mouse anti-PSD-95 (1:400, ab2723, Abcam), mouse anti-GFP (1:5000, ab290, Abcam), rabbit anti-Homer1a/b/c (1:1000, 160-002, Synaptic Systems), mouse anti-Bassoon (1:400, 141-011, Synaptic Systems), guinea pig anti-MAP2 (1:500, 188-004, Synaptic Systems), mouse anti-ubiquitin (1:1000, BML-PW8810-0100, Enzo), mouse anti-γ−tubulin (1:10 000, T7451, Sigma-Aldrich), mouse anti-HA (1:5000, H3663, Sigma-Aldrich). Secondary antibodies used were IgG– horseradish peroxidase goat (1:3000; sc-2768; Santa Cruz Biotechnology), mouse (1:3000; sc-2380; Santa Cruz Biotechnology), and rabbit (1:3000; 31466; Thermo Fisher Scientific). Alexa Fluor 488 anti–rabbit, Alexa Fluor 594 anti–mouse, Alexa Fluor 647 anti–guinea pig, (1:1,000, A-11034, A-11032, A-21450, Thermo Fisher Scientific). Picrotoxin, tetrodotoxin and Nimodipine were from Tocris, WP1130 was from APExBIO and Lactacystin was from Santa Cruz Biotechnology.

Dissociated cultures of primary hippocampal neurons were prepared from E18 Sprague-Dawley rat embryos as described previously (Richier et al., 2010). All animal experiments and procedures were approved by the Dalhousie University Committee on Laboratory Animals (protocol no. 23-072) and conducted in accordance with the guidelines of the Canadian Council on Animal Care (CCAC). HEK293 and IEC-18 cells were maintained at 37°C and 5% CO2 in DMEM supplemented with 10% FBS or 5% FBS (IEC-18) and 10 U/ml penicillin and 10 μg/ml streptomycin. HEK293 cells were transfected with polyethylenimine (PEI) by mixing cDNA with PEI (1 mg/ml) at a ratio of 4:1 PEI to DNA. For transfection of neurons with cDNA, the calcium phosphate method was used as previously described (Richier et al., 2010). For AMOT-130 knockdown in IEC-18 cells, 20 µM siRNA was transfected to a final concentration of 10 nM using Lipofectamine 2000 (Thermo Fisher Scientific) according to the manufacturer’s recommendations. Cells were harvested 2 days later and the expression levels of endogenous AMOT-130 was analyzed by immunoblotting using C-18 antibody. For biochemical analysis, neurons were solubilized in lysis buffer containing: (20 mM Tris [pH 8.0], 137 mM NaCl, 10 % glycerol, 1 % NP-40), supplemented with protease inhibitors and lysates were centrifuged at 13 800 g for 20 minutes at 4°C. Supernatant fractions were analyzed by Western blotting.

For knockdown of AMOT-130 in neurons, DIV7 cultures were transfected with siRNA together with 20 µg pEGFP plasmid using Lipofectamine 2000 and prepared for imaging at DIV9. To induce neuronal activity, DIV11 hippocampal neurons were treated with vehicle (DMSO, v/v), picrotoxin (50 µM), or tetrodotoxin (1 µM) for 24 hours in minimum essential medium (MEM) supplemented with B27 (Thermo Fisher). Neurons were treated with WP1130 (4 µM) alone or together with Lactacystin (100 µM) for 4 hours. DIV7 cultures were pretreated with Nimodipine (5 µM), an L-type calcium channel blocker, for 30 minutes followed by either high KCl (50 mM) or control media containing NaCl (50 µM) for another 40 minutes. Neurons were fixed in 4% PFA with 4% sucrose in 1x PBS for 10–20 min at RT and permeabilized in 0.1% Triton X-100 for 10 min. Fixed cells were washed extensively in 1x PBS and then blocked in 5% BSA in PBS with 0.1% Triton X-100 for 1 h. After blocking, cells were incubated with primary antibodies in 5% BSA overnight at RT. Then, cells were washed in 1x PBS three times for 2 minutes each, incubated with secondary antibodies in 5% BSA for 1 hour at RT, washed in 1x PBS, and mounted on microscope slides with fluoromount (Sigma-Aldrich).

For immunoprecipitation in primary hippocampal cultures, crude synaptosomal P2 fractions prepared as previously described (Wigerius et al., 2018), were solubilized at RT in lysis buffer and centrifuged at 13 800 g for 45 minutes at 4°C. Lysates were incubated with 1–2 µg of either isotype-matched IgG, αNedd4-1 or α2095 antibody at 4°C overnight followed by incubation with protein-A Sepharose beads (GE Healthcare).

Protein concentrations were measured using the bicinchoninic assay method (Thermo Fisher Scientific). Bound proteins were subjected to SDS-PAGE and detected by Western blotting using the indicated antibodies. Relative protein intensities were quantified by densitometry using Fiji (Image J, National Institutes of Health) software.

The ubiquitylation assay was performed under denaturing conditions to ensure complete dissociation of AMOT binding proteins as follows: neurons plated onto 10 cm plates were treated with PTX for 24 hours, then harvested and washed in ice-cold 1xPBS. To assure ubiquitylation of endogenous AMOT and not associated proteins, a cytosolic protein fraction was prepared in 200 µl of ubiquitin lysis buffer (20 mM Tris/HCl, pH 7.5, 300 mM NaCl and 1 % Triton X-100 supplemented with protease inhibitors) from which 40 µl was saved for Western blotting. To the remaining cytosolic fraction of 160 µl, 16 µl of 10 % SDS was added to a final concentration of 1 %. Samples were boiled twice for 5 minutes combined with vigorous mixing. After adding 704 µl of ubiquitin buffer to reduce the SDS concentration to 0.2 % lysates were collected by centrifugation at 13,800 g for 15 minutes. Lysates were then incubated with 2 µg of α2095 antibody overnight at 4°C, followed by incubation with protein-A Sepharose beads (GE Healthcare). Beads were washed three times in ubiquitin lysis buffer, eluted in 4xLaemmli sample buffer and prepared for SDS-PAGE analysis.

Images were acquired as z-stack series taken at 0.3-μm step intervals with an LSM710 confocal scanning microscope using 40x 0.7 NA and 63x 1.4 NA oil immersion lenses and collected with a digital Axiocam camera controlled by ZEN software (ZEISS). The settings and acquisition parameters were the same across all experiments. Data was collected from at least two independent experiments per condition and analyzed with the experimenter blinded to the conditions. To analyze dendrite growth z-stacks of 5-10 images were converted into two-dimensional images by maximum intensity projection. Tracing and quantification of total dendritic branch tip number (TDBTN) and total dendritic branch length (TDBL) was performed with Fiji using the Neuron J plugin (Meijering et al., 2004). To analyze AMOT-130 punctate after pharmacological treatment, z-stacks were converted by maximum-intensity projection. The threshold for AMOT-130 was adjusted and chosen such that all punctate included in the analysis were overlapping or apposing bassoon punctate within a distance of 0.5 µm from the center of the cluster. Binary masks with pixel intensities were created and the punctate were highlighted by generating ROI masks that were applied along 100-µm dendritic segments labeled with MAP2. The values were presented as total fluorescence of endogenous AMOT-130 punctate before and after treatment.

Statistical analysis was performed in Prism (6.0) software, and all datasets were checked for Gaussian normality distribution. Data testing of two groups was performed by a two-tailed unpaired Student’s t test. Multiple group analysis was performed using one-way ANOVA followed by the Tukey-Kramer post hoc test to account for multiple comparisons. Statistical significance was set at P < 0.05. All figures were created in Illustrator CS5 (Adobe).

## Results

Synapse and dendrite stabilization are strongly coupled during early development, whereas this dependence diminishes as neurons mature as a result of synaptic pruning (Koleske, 2013). AMOT-130 regulates dendritic spine development(Rojek et al., 2019), and loss of all AMOT isoforms affects dendritic complexity (Rojek et al., 2019).

However, whether AMOT-130 or AMOT-80 contributes to this remains unknown. To test for this possibility, we assessed the impact of silencing AMOT-130 alone on dendrite morphology in young hippocampal cultures using RNAi (**Fig 1A**). The efficacy of control (CsiRNA) and small-interfering (si) RNAs (siRNA#1 and siRNA#2) were tested in rat IEC-18 cells due to high transfection efficiency. Both siRNAs reduced the levels of endogenous AMOT-130 by at least 86% compared to cells transfected with the control (**Fig 1B**). We next introduced the siRNA in DIV7 hippocampal neurons together with plasmid encoding Green fluorescent protein (GFP) and imaged cells 2 days later (**Fig 1C**). To evaluate neuronal morphology of reconstructed branches (**Fig 1C, bottom panels**), we quantified the total dendritic branch tip number (TDBTN) and total dendritic branch length (TDBL) as measures of dendritic arbor development (Liao, Bird, Cuntz, & Howard, 2023). Silencing of AMOT-130 reduced TDBTN by 38% (17.1 ± 3.50) and 33% (18.5 ± 5.24) following treatment with siRNA#1 and siRNA#2, respectively, compared with control (27.5 ± 1.50; Fig 1D; **** P < 0.00001, one-way ANOVA with Tukey’s multiple-comparison test). Likewise, TDBL was reduced by 54% (604.2 ± 54.4 µm) and 52% (631.1 ± 69.1 µm), respectively, relative to control (1311 ± 104.3 µm; **Fig 1E**; **** P < 0.00001, one-way ANOVA with Tukey’s multiple-comparison test). Moreover, changes in TDBTN and TDBL were tightly correlated for all siRNA treated neurons (**Fig 1F**; r = 0.630; **** P < 0.00001, Pearson correlation analysis), consistent with an association between reduced dendritic branching and length. Taken together, these results extend the finding by Rojek (Rojek et al., 2019), showing that it is specifically the AMOT-130 isoform that regulates dendritic complexity, mirroring the role of this isoform in the structural plasticity of dendritic spines in mature neurons.

**Figure 1.**
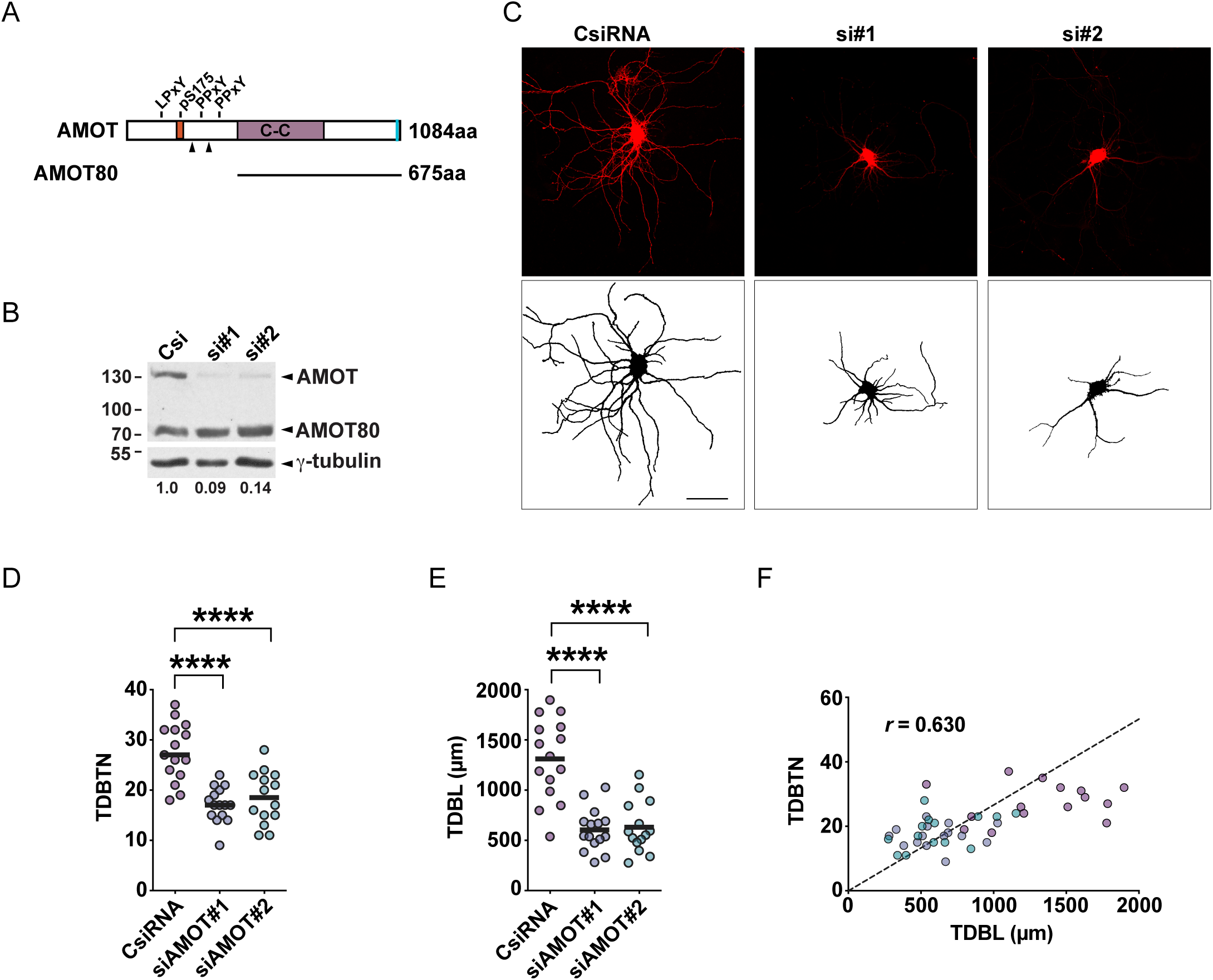
AMOT-130-specific knockdown reduces dendritic complexity in hippocampal neurons. **A.** Domain structure of AMOT-130. The positions of the L/PPxY motifs and the actin-binding motif (orange) are indicated. The C-terminal region comprising a coiled-coil domain and a PDZ-binding motif (turquoise) are identical to the corresponding regions in the AMOT-80 isoform. Arrowheads indicate the RNAi target sites in the N-terminal region specific to the AMOT-130 isoform. **B.** Rat IEC-18 cell lysate transfected with control (CsiRNA) or AMOT-130 siRNA (siRNA#1 and siRNA#2) was analyzed by immunoblotting with antibody recognizing both AMOT-130 and AMOT-80, and re-probed against γ-tubulin. Numbers indicate residual amounts of AMOT-130 relative to control. **C.** Rat hippocampal neurons were co-transfected with plasmid encoding pEGFP (pseudo-colored) and AMOT-130 siRNA on day in vitro (DIV) 7 and imaged on DIV9. Scalebar, 50µm. **D.** Plot of Total Dendritic Branch Tip Number (TDBTN) in AMOT-130 siRNA-treated neurons. n > 14 neurons in each condition. TDBTN: CsiRNA, 27.5 ± 1.50 versus siRNA#1, 17.1 ± 3.50 and siRNA#2, 18.5 ± 5.24. **** P < 0.00001, one-way ANOVA with Tukey’s multiple-comparison test. **E.** Plot of Total Dendritic Branch Length (TDBL) in AMOT-130 siRNA-treated neurons. CsiRNA, 1311 ± 104.3 µm versus siRNA#1, 604.2 ± 54.4 µm and siRNA#2, 631.1 ± 69.1 µm. **** P < 0.00001, one-way ANOVA with Tukey’s multiple-comparison test. **F.** Pearson correlation plot of TDBTN versus TDBL, with linear regression indicated by the dotted line: n > 42 neurons for all conditions. r = 0.6303. **** P < 0.00001; Pearson correlation analysis.

Global changes in synaptic activity levels have previously been shown to lead to homeostatic changes in the expression levels of synaptic proteins [Ehlers, 2003]. Since AMOT is localized to synaptic spines and regulated in its localization by phosphorylation (Wigerius et al., 2018), we asked whether asked whether AMOT-130 expression is altered in response to neuronal activity. To test this idea, we treated cultures for 24 hours with either picrotoxin (PTX, 50 µM), a GABA-R blocker that increases excitatory synaptic activity, or tetrodotoxin (TTX, 1 µM), which blocks action potential-dependent synaptic activity (Redmond, Kashani, & Ghosh, 2002; Turrigiano, Leslie, Desai, Rutherford, & Nelson, 1998). Following treatment, neurons were immunostained for endogenous AMOT-130 together with the dendritic marker MAP2 and the excitatory presynaptic marker Bassoon (**Fig 2A**). Treatment with PTX revealed a marked reduction of AMOT-130 fluorescence puncta intensity by more than 65% compared with vehicle or TTX (**Fig 2B**; **** P < 0.00001; one-way Anova with Tukey’s multiple comparisons test), suggesting that AMOT-130 levels are decreased in response to strongly elevated activity.

**Figure 2.**
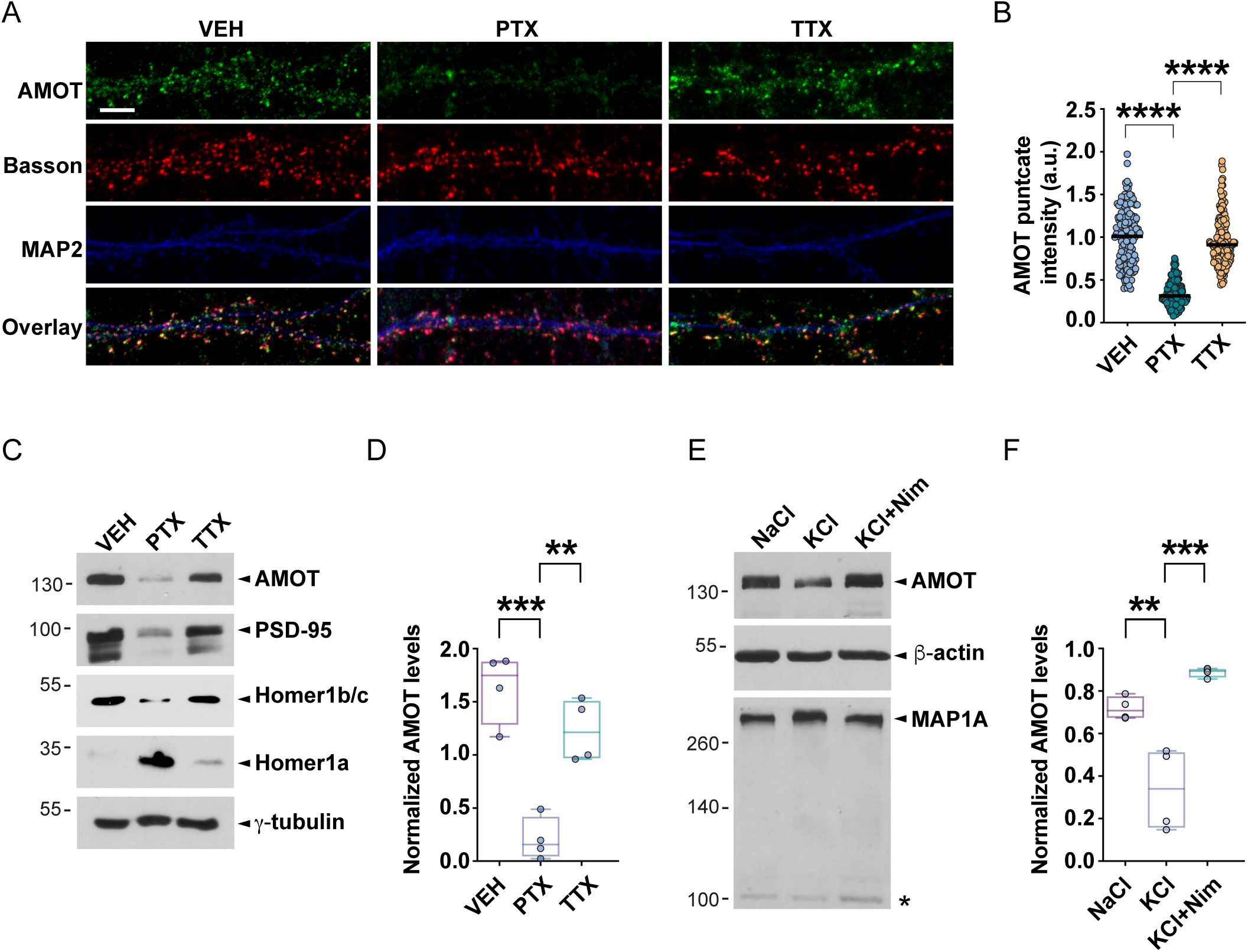
Neuronal excitation reduces synaptic AMOT-130 punctate protein levels in hippocampal neurons. **A.** DIV9 hippocampal neurons treated with Vehicle (VEH), 50 µM picrotoxin (PTX), or 1 µM TTX for 24 hours were fixed and coimmunostained with antibodies that recognizes AMOT-130, Bassoon and MAP2. Scale bar, 5 µM. **B.** The box plot shows changes in AMOT-130 punctate intensity along dendrite segment. n = 100 punctate in each condition. **** P < 0.00001; one-way ANOVA with Tukey’s multiple-comparison test. **C.** Lysates from DIV10 hippocampal cultures treated with vehicle, PTX and tetrodotoxin, TTX, (1 µM) for 24 hours were analyzed by immunoblotting with the indicated antibodies. **D.** Box plot showing AMOT-130 protein levels normalized to tubulin. n = 4 independent experiments. ***, P < 0.0001, **, P < 0.001, one-way ANOVA with Tukey’s multiple-comparison test. **E.** Lysates from DIV7 hippocampal cultures pretreated with Nimodipine, Nim, (5 µM) followed by high KCl (50 mM), or NaCl (50 mM, control) for 1 hour were analyzed by immunoblotting with the indicated antibodies. **F.** Box plot showing AMOT-130 protein levels normalized to actin. n= 4 independent experiments. ***, P < 0.0001, **, P < 0.001, one-way ANOVA with Tukey’s multiple-comparison test.

To confirm the loss of AMOT-130 expression, we analyzed whole cell lysate from activated and inactivated neurons. Activity reduced the relative AMOT-130 expression by approximately 8-fold relative to untreated neurons, whereas blockade of synaptic activity retained levels similar to untreated controls (**Fig 2C, D**; vehicle relative to PTX, *** P < 0.0001; PTX relative to TTX, **P < 0.001; vehicle relative to TTX, P = 0.156; one-way Anova with Tukey’s multiple comparisons test). In addition, we detected a similar reduction in the levels of PSD-95 and Homer 1b/c, but not Homer 1a, as previously reported (Hu et al., 2010; Piccoli et al., 2007; Sun & Turrigiano, 2011). Importantly, γ-tubulin expression remained unchanged across treatments confirming that the observed changes were specific to the treatment and proteins examined (**Fig 2A**). Since different activity protocols produce distinct outcomes on protein expression (Lukin et al., 2025), we also tested whether potassium-induced neuronal depolarization, a process dependent on L-type calcium-channel activation, influence AMOT-130 expression. DIV 7 neurons were stimulated with extracellular KCl followed by immunoblot analysis. As a positive control for KCl-induced depolarization we monitored MAP1A, which has previously been shown to be up-regulated in response to high KCl (Szebenyi et al., 2005). A 60-min treatment with KCl (50 mM) consistently reduced AMOT-130 expression and increased MAP1A levels compared to NaCl (50 mM) while treatment with Nimodipine (5 µM) an L-type calcium-channel blocker restored both AMOT-130 and MAP1A levels (**Fig 2E**, F; NaCl relative to KCl, *** P < 0.0001; KCl versus KCl + Nim, ** P < 0.001; one-way Anova with Tukey’s multiple comparisons test). Taken together, these results provide evidence that AMOT-130 protein abundance decreases in response to increased excitatory neuronal activity induced by either PTX or KCl in cultured neurons.

Activity-dependent changes in AMOT-130 expression could result from altered protein synthesis or increased protein degradation, as reported for several postsynaptic proteins following elevated activity (Schanzenbacher, Sambandan, Langer, & Schuman, 2016). Activity-dependent changes in synaptic strength have also been associated with increased levels of Nedd4-1, a member of the Nedd4 family of ubiquitin ligases that target some angiomotin proteins for degradation (Scudder et al., 2014; C. Wang et al., 2012). These observations suggested that the UPS might mediate downregulation of AMOT-130 in response to elevated activity. Additionally, AMOT-80 has been suggested to be a target for ubiquitin-mediated degradation (C. Wang et al., 2012). We therefore initially examined the Nedd4-1 interaction in HEK293 lysate using an antibody that recognizes both the AMOT-130 and 80 isoforms. Nedd4-1 co-immunoprecipitated AMOT-130 but not the shorter isoform AMOT-80, indicating that it is not a target of endogenous Nedd4-1 (**Fig 3A**). To confirm the previously reported involvement of the three N-terminal L/PxYY motifs for AMOT-130 binding (C. Wang et al., 2012), we expressed YFP-AMOT-AAA, in which the three tyrosine residues were replaced with alanine (Fig 1A). Consistent with earlier findings, the mutant showed markedly reduced association with Myc-Nedd4-1 (**Fig 3B**). We next examined whether endogenous Nedd4-1 and AMOT-130 associate in hippocampal neurons by immunoprecipitating endogenous AMOT-130 from hippocampal lysate using an isoform specific antibody (# 2095), followed by immunoblotting for Nedd4-1. Nedd4-1 was specifically detected in the AMOT-130 precipitates (**Fig 3C**) confirming that they associate in neurons. The link between AMOT-130 with Nedd4-1, suggests that AMOT-130 synaptic levels are counterbalanced by the activity of the UPS and DUBs to maintain homeostasis. To test this idea, we treated hippocampal neurons with WP1130 (4 µM) alone or together with the proteasome specific inhibitor Lactacystin (100 µM) for 4 hours. WP1130 is a selective DUB inhibitor that has been described to shift AMOT-130 stability towards ubiquitin-mediated degradation (Thanh Nguyen et al., 2016). While treatment with WP1130 resulted in a significant reduction of AMOT-130 levels, simultaneous blockage of both the proteasome and DUBs restored AMOT-130 levels in neurons (**Fig 3D, E**; Vehicle relative to WP1130, *** P < 0.0001; WP1130 relative to Lact + WP, * P < 0.05; one-way Anova with Tukey’s multiple comparisons test). Taken together, these data suggest that AMOT-130 expression in neurons is bi-directionally controlled by the UPS and DUBs. To determine whether elevated neuronal activity promotes ubiquitination, we immunoprecipitated endogenous AMOT-130 from neurons treated with WP1130 or PTX using the α2095 antibody. Immunoprecipitates were then probed with an antibody recognizing conjugated polyubiquitin chains. Both treatments markedly increased AMOT-130 polyubiquitination compared with vehicle-treated neurons (**Fig 3F**). These findings support the idea that neuronal activity regulates AMOT-130 expression levels through the UPS, supporting a recent study showing the importance of USP9x in association with WWC1/2 in the regulation of AMOT stability(Cao et al., 2023).

**Figure 3.**
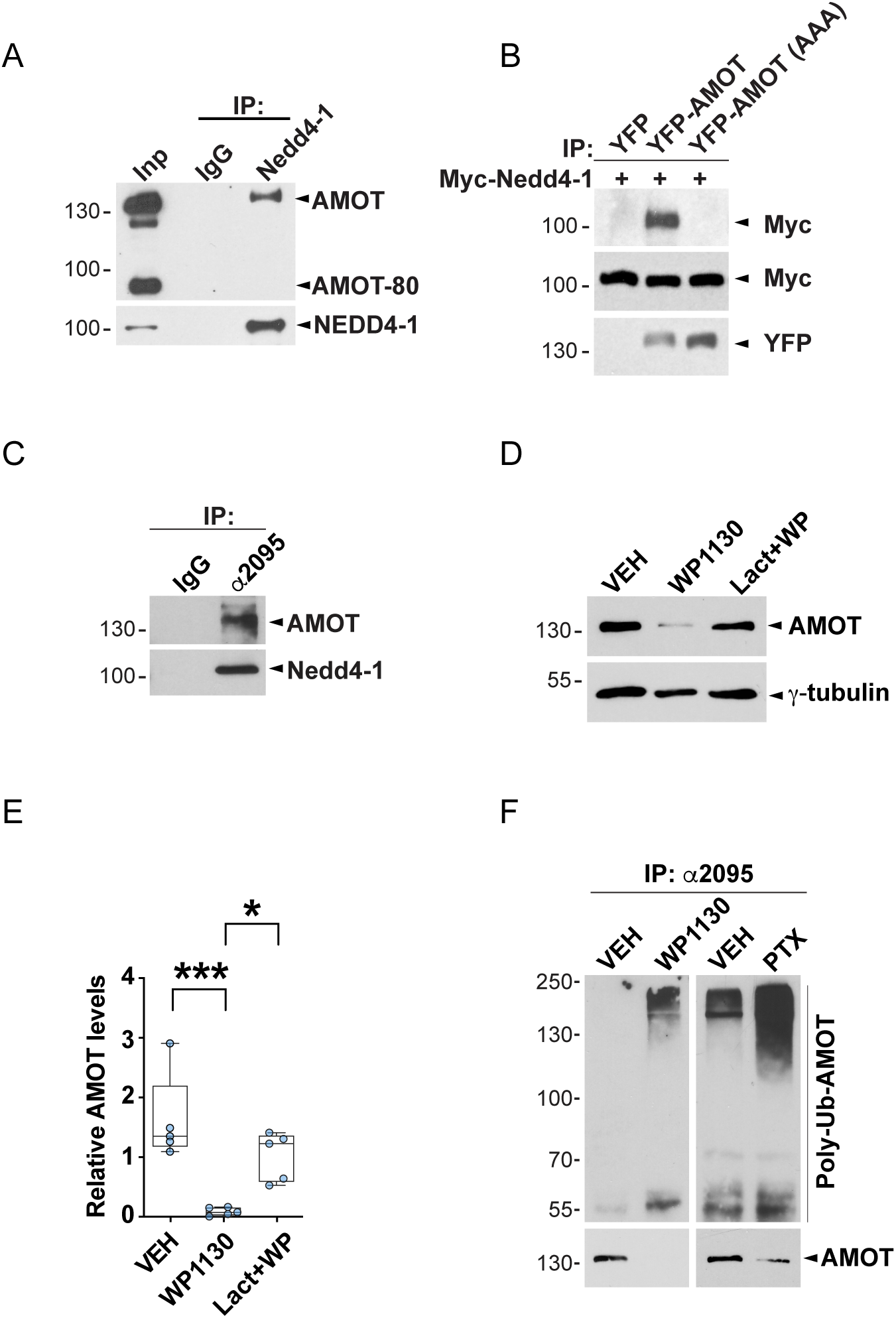
AMOT-130 associates with Nedd4-1 and undergoes activity-dependent proteasomal degradation in hippocampal neurons. **A.** Lysates from HEK293 cells immunoprecipitated (IP) with antibody recognizing control IgG or endogenous Nedd4-1 were analyzed by immunoblotting with antibody recognizing both AMOT isoforms and re-probed with anti-Nedd4-1 antibody. Input (Inp). **B.** Lysates from HEK293 cells co-transfected with plasmid encoding Myc-Nedd4-1 together with either YFP, YFP-AMOT or YFP-AMOT-AAA (all three L/PPxY motifs mutated to alanine), were immunoprecipitated with anti-GFP antibody. Bound proteins were analyzed by immunoblotting with anti-Myc and anti-GFP antibodies. **C.** Lysates from DIV15 hippocampal cultures immunoprecipitated with IgG or antibody (α2095) that recognizes the unique N-terminal region in AMOT were analyzed by immunoblotting. Bound proteins were detected with anti-Nedd4-1 antibody and re-probed with the α2095 antibody. **D.** Lysates from hippocampal cultures treated with vehicle, WP1130 (4 µM) alone or together with Lactacystin (Lact, 100 µM) for 4 hours were analyzed by immunoblotting with α2095 antibody and re-probed with γ−tubulin antibody. Box plot (bottom) shows AMOT-130 protein normalized to γ-tubulin. n = 3 independent experiments. ***, P < 0.0001, *, P < 0.05, one-way ANOVA with Tukey’s multiple-comparison test. **D.** Lysates from hippocampal cultures treated with vehicle, WP1130 for 4 hours or PTX for 24 hours were immunoprecipitated with α2095 antibody and analyzed by immunoblotting. AMOT-130 was immunoprecipitated using the α2095 antibody and associated polyubiquitin conjugates were detected using an anti-Ub antibody. AMOT-130 input (bottom) was 10% of total lysate.

Given that activity affects the stability of AMOT-130 and its ubiquitination, we next wanted to determine the effects of PTX treatment on neuronal morphology. Reconstruction of dendritic arbors revealed reduced complexity in response to synaptic activity., Interestingly, the over-expression of AMOT-130 protected against this loss (**Fig 4A**). Quantification of dendrites demonstrated that activity reduced TBTN by 21% (**Fig 4D**; 16.2 ± 0.75 relative to 12.8 ± 0.62; ** P < 0.001; one-way Anova with Tukey’s multiple comparisons test) and TDBL by 26 % (**Fig 4E**; 1632 ± 87.5 µm relative to 1207 ± 72.4 µm; ** P < 0.001; one-way Anova with Tukey’s multiple comparisons test) whereas the same treatment in neurons overexpressing AMOT-130 did not reduce either TDBTN or TDBL (**Fig 4D, E**). Consistent with these findings, plotting TDBTN against TDBL for all neurons showed a linear relationship across the experimental conditions (**Fig 4F**; r = 0.475; *** P < 0.0001, Pearson correlation analysis).

**Figure 4.**
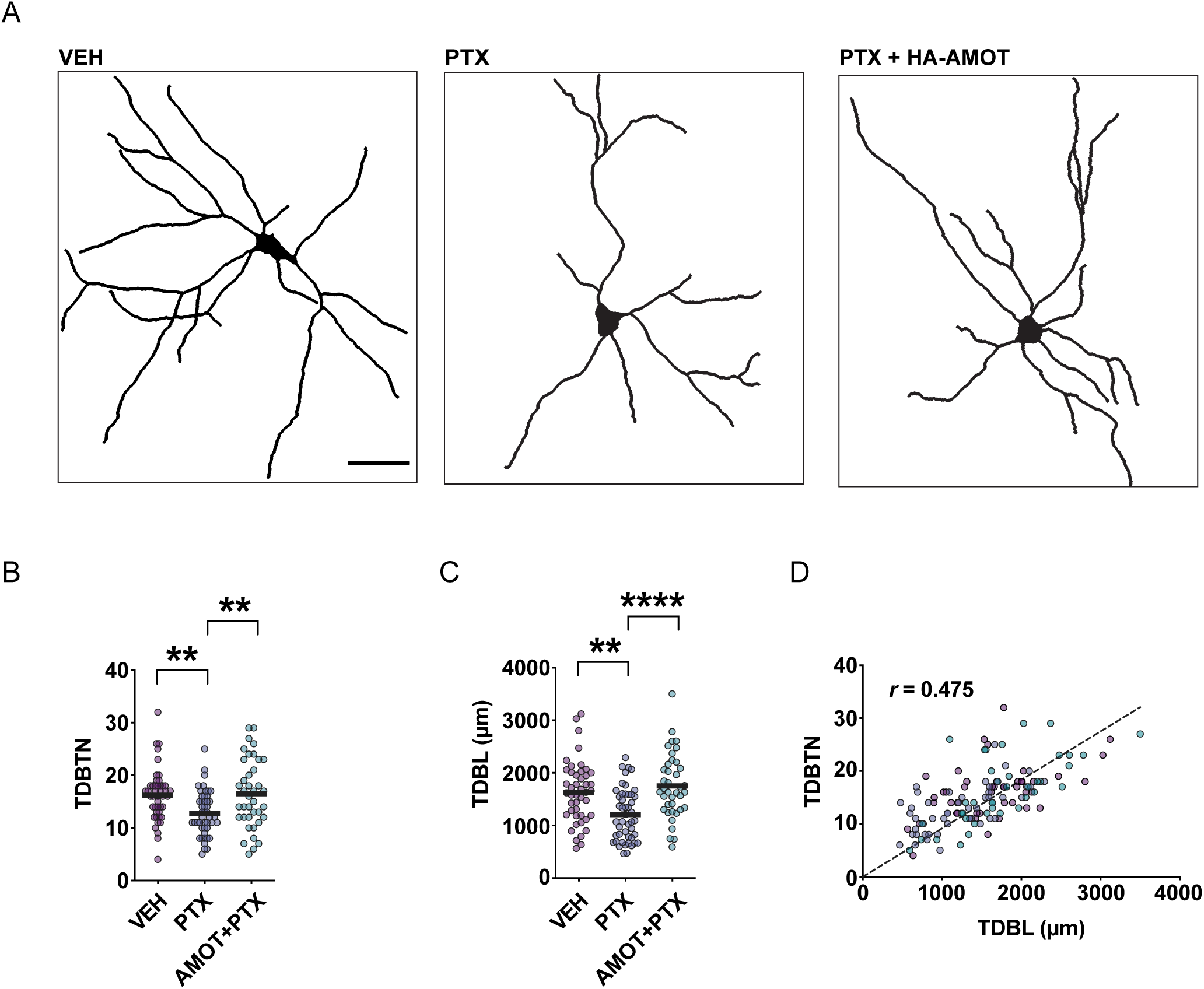
Neuronal excitation reduces neuronal complexity which is rescued by AMOT-130 expression. **A.** 2D-reconstruction of neurons imaged at DIV10 expressing pEGFP alone or together with HA-AMOT-130, and treated with vehicle, PTX or TTX for 24 hours. Scale bar, 20 µm. **B.** Plot of TDBTN in neurons expressing vehicle, 16.2 ± 0.75 versus PTX, 12.8 ± 0.62 and PTX+AMOT-130, 16.5 ± 1.04. n > 39 neurons in each condition. ** P < 0.001, one-way ANOVA with Tukey’s multiple-comparison test. **C.** Plot of TDBL in neurons expressing vehicle, 1632 ± 87.5 µm versus PTX, 1207 ± 72.4 µm and PTX+AMOT-130, 1750 ± 98.3 µm. **, P < 0.001, ****, P < 0.00001, one-way ANOVA with Tukey’s multiple-comparison test. **D.** Pearson correlation plots of TDBTN versus TDBL, with linear regression indicated by the dotted line: n > 117 neurons for all conditions. r = 0.475. ***, P < 0.0001; Pearson correlation analysis.

## Discussion

We previously showed that AMOT-130 regulate dendritic spine morphogenesis in primary hippocampal neurons (Wigerius et al., 2018) while others showed reduced dendritic complexity of cortical neurons (Rojek et al., 2019). However, the study by Rojek et al used a combined knockdown of AMOT-130 and AMOT-80 leaving their individual contribution to dendritic arborization unresolved. To address this, we selectively depleted AMOT-130 and found that its loss alone reduces dendritic arbor complexity. Moreover, we noted that AMOT-130 levels in the rodent brain steadily increased during neuronal maturation (Wigerius et al., 2018) and since activity drives neuronal growth during this developmental period, we asked whether AMOT-130 responds to different activity paradigms *in vitro*. He we report that elevated activity downregulates AMOT-130 through the ubiquitin-proteasome system. Intriguingly, this downregulation appears to be required for the modulation of dendritic remodeling by activity, as overexpression of AMOT130 prevented PTX-mediated reduction of dendritic complexity, a phenotype also observed in neurons silenced for AMOT-130. Together, these results define synaptic scaling as a control mechanism that regulates dendritic remodeling by modulating AMOT-130 protein levels in primary hippocampal neurons.

We detected reduced Homer 1b/c and PSD-95 protein levels, consistent with their reported downscaling in response to elevated activity (Sun & Turrigiano, 2011; Swann, Le, Lam, Owens, & Mayer, 2007). However, unlike Homer and PSD-95, this is the first report linking AMOT-130 to homeostatic synaptic scaling suggesting that these proteins belong to the same co-regulated synaptic cluster. Moreover, loss of either AMOT-130 and PSD-95/Homer 1b/c results in spine regression through the formation of filopodia-like intermediates. It is therefore likely that they are also coupled to one another in functional ensembles that share organizational similarities. Homer 1a, in contrast, exhibited increased expression, reflecting its role in triggering synaptic downscaling leading to structural remodeling of spines (Diering et al., 2017; Hu et al., 2010). We further discovered that high extracellular KCl mimicked the effect of PTX on AMOT-130 levels, whereas MAP1A, a marker of dendritic branching and synapse growth, displayed enhanced expression as previously reported (Szebenyi et al., 2005). Blockage of L-type calcium channels prevented the down regulation, suggesting that calcium influx is required for the reduction in AMOT-130 expression. Notably, calcium enhances Lats1 kinase activity (Thines, Gorisse, Li, Sayedyahossein, & Sacks, 2022), and we have demonstrated that Lats1-dependent phosphorylation of AMOT-130 restricts its enrichment in spines. Thus, whether Lats1 also contributes to the scaling down effect by priming AMOT-130 for subsequent downregulation remains to be determined.

UPS mediated degradation has previously been reported to regulate the abundance of synaptic proteins in response to activity (Yi & Ehlers, 2005). We here propose that AMOT-130 levels in neurons are likewise controlled by the activity of the UPS and DUB enzymes. First, we find that Nedd4-1, a E3 ligase which has been shown to increase its expression in response to elevated activity (Scudder et al., 2014), associates with AMOT-130 in the rodent brain, and that the N-terminal PPxY motifs are necessary for this association confirming other studies in non-neuronal cells (C. Wang et al., 2012). While several family members have been reported to target AMOT-130 in different contexts, we note that Nedd4-1 is, to our knowledge, the only member that has been reported to regulate dendritic remodeling in hippocampal neurons (Kawabe et al., 2010). Second, AMOT-130 degradation in our study was counteracted by the activity of DUBs which are highly expressed in the developing brain (Todi & Paulson, 2011). Importantly, USP9x, a DUB previously found to specifically counteract NEDD4-1-mediated AMOT-130 degradation, has also been implicated in maintaining protein abundance at the PSD (Kwon et al., 2016; Yoon, Parnell, & Penzes, 2020). Further, WWC1 has recently been shown to recruit USP9x to affect the stability of AMOT and contribute to cognition, whether this is linked to activity-dependent AMOT-130 regulation remains to be determined (Cao et al., 2023). Third, pharmacological blockage of either the UPS or DUBs prevents both ubiquitin conjugation and deconjugation, suggesting that the ubiquitination state of AMOT-130 is actively maintained in neurons. We conclude therefore that homeostatic scaling provides a selective mechanism for counterbalancing the elevated expression levels of AMOT-130 during synaptic development.

In summary, our data suggests that AMOT-130 expression is reversibly regulated by synaptic activity and involves the UPS which influences dendritic arbors. It is likely that other mechanism that mediate changes in synaptic strength are involved. Yet, given that synapse and dendrite stabilization are coupled during development, synaptic scaling of AMOT-130 at all synapses might contribute to the observed reduction in dendritic complexity. A key goal in future will be to identify whether degradation of AMOT-130 results in synaptic weakening as predicted by our study. It will also be important to define the molecular networks surrounding AMOT-130 in these contexts including actin-binding proteins and components of the Hippo pathway and their targets. Finally, as Motin family of proteins can homo- and hetero-oligomerize through their coiled-coiled domain (Ernkvist et al., 2008; Patrie, 2005), it will be interesting to determine the function of AMOT-80 in post-mitotic neurons and its contribution to morphology and activity dependent mechanisms.

## Acknowledgements

This work was supported by funding from Discovery grants RGPIN-2022-03370 (J.P.F) and RGPIN-2021-03303 (S.R.K) from the National Science and Engineering Council (NSERC) of Canada. The authors declare no conflict of interest.

## References

Ambrozkiewicz, M. C., & Kawabe, H. (2015). HECT-type E3 ubiquitin ligases in nerve cell development and synapse physiology. FEBS Lett, 589(14), 1635–1643. doi:10.1016/j.febslet.2015.05.009

Bingol, B., & Schuman, E. M. (2006). Activity-dependent dynamics and sequestration of proteasomes in dendritic spines. Nature, 441(7097), 1144–1148. doi:10.1038/nature04769

Cao, R., Zhu, R., Sha, Z., Qi, S., Zhong, Z., Zheng, F.,…Yu, F. X. (2023). WWC1/2 regulate spinogenesis and cognition in mice by stabilizing AMOT. Cell Death Dis, 14(8), 491. doi:10.1038/s41419-023-06020-7

Cingolani, L. A., & Goda, Y. (2008). Actin in action: the interplay between the actin cytoskeleton and synaptic efficacy. Nat Rev Neurosci, 9(5), 344–356. doi:10.1038/nrn2373

Diering, G. H., Nirujogi, R. S., Roth, R. H., Worley, P. F., Pandey, A., & Huganir, R. L. (2017). Homer1a drives homeostatic scaling-down of excitatory synapses during sleep. Science, 355(6324), 511–515. doi:10.1126/science.aai8355

Dogterom, M., & Koenderink, G. H. (2019). Actin-microtubule crosstalk in cell biology. Nat Rev Mol Cell Biol, 20(1), 38–54. doi:10.1038/s41580-018-0067-1

Ehlers, M. D. (2003). Activity level controls postsynaptic composition and signaling via the ubiquitin-proteasome system. Nat Neurosci, 6(3), 231–242. doi:10.1038/nn1013

Ernkvist, M., Birot, O., Sinha, I., Veitonmaki, N., Nystrom, S., Aase, K., & Holmgren, L. (2008). Differential roles of p80- and p130-angiomotin in the switch between migration and stabilization of endothelial cells. Biochim Biophys Acta, 1783(3), 429–437. doi:10.1016/j.bbamcr.2007.11.018

Falace, A., Nosrati, M. S. S., Dostmohammadi, A., Doustmohammadi, M., Striano, P., Zara, F.,…Scala, M. (2026). Wiring the developing brain: The Rho GTPase signaling network in neurodevelopmental disorders. Neurobiol Dis, 229, 107609. doi:10.1016/j.nbd.2026.107609

Hu, J. H., Park, J. M., Park, S., Xiao, B., Dehoff, M. H., Kim, S.,…Worley, P. F. (2010). Homeostatic scaling requires group I mGluR activation mediated by Homer1a. Neuron, 68(6), 1128–1142. doi:10.1016/j.neuron.2010.11.008

Kawabe, H., Neeb, A., Dimova, K., Young, S. M., Jr., Takeda, M., Katsurabayashi, S.,…Brose, N. (2010). Regulation of Rap2A by the ubiquitin ligase Nedd4-1 controls neurite development. Neuron, 65(3), 358–372. doi:10.1016/j.neuron.2010.01.007

Koleske, A. J. (2013). Molecular mechanisms of dendrite stability. Nat Rev Neurosci, 14(8), 536–550. doi:10.1038/nrn3486

Kwon, S. C., Nguyen, T. A., Choi, Y. G., Jo, M. H., Hohng, S., Kim, V. N., & Woo, J. S. (2016). Structure of Human DROSHA. Cell, 164(1-2), 81–90. doi:10.1016/j.cell.2015.12.019

Lefebvre, J. L., Sanes, J. R., & Kay, J. N. (2015). Development of dendritic form and function. Annu Rev Cell Dev Biol, 31, 741–777. doi:10.1146/annurev-cellbio-100913-013020

Liao, M., Bird, A. D., Cuntz, H., & Howard, J. (2023). Topology recapitulates morphogenesis of neuronal dendrites. Cell Rep, 42(11), 113268. doi:10.1016/j.celrep.2023.113268

Lukin, J., Beckel, M. S., Pedroncini, O., Giusti, S. A., Di Mauro, G. C., Linenberg, I. M.,…Refojo, D. (2025). Neuronal activity-dependent gene expression is stimulus-specific and changes with neuronal maturation. Front Mol Neurosci, 18, 1609772. doi:10.3389/fnmol.2025.1609772

Matsuzaki, M., Honkura, N., Ellis-Davies, G. C., & Kasai, H. (2004). Structural basis of long-term potentiation in single dendritic spines. Nature, 429(6993), 761–766. doi:10.1038/nature02617

Matt, L., Kim, K., Chowdhury, D., & Hell, J. W. (2019). Role of Palmitoylation of Postsynaptic Proteins in Promoting Synaptic Plasticity. Front Mol Neurosci, 12, 8. doi:10.3389/fnmol.2019.00008

Matus, A. (2000). Actin-based plasticity in dendritic spines. Science, 290(5492), 754–758. doi:10.1126/science.290.5492.754

Meijering, E., Jacob, M., Sarria, J. C., Steiner, P., Hirling, H., & Unser, M. (2004). Design and validation of a tool for neurite tracing and analysis in fluorescence microscopy images. Cytometry A, 58(2), 167–176. doi:10.1002/cyto.a.20022

Murakoshi, H., Wang, H., & Yasuda, R. (2011). Local, persistent activation of Rho GTPases during plasticity of single dendritic spines. Nature, 472(7341), 100–104. doi:10.1038/nature09823

Nakahata, Y., & Yasuda, R. (2018). Plasticity of Spine Structure: Local Signaling, Translation and Cytoskeletal Reorganization. Front Synaptic Neurosci, 10, 29. doi:10.3389/fnsyn.2018.00029

Okamoto, K., Nagai, T., Miyawaki, A., & Hayashi, Y. (2004). Rapid and persistent modulation of actin dynamics regulates postsynaptic reorganization underlying bidirectional plasticity. Nat Neurosci, 7(10), 1104–1112. doi:10.1038/nn1311

Pan, Y., & Monje, M. (2020). Activity Shapes Neural Circuit Form and Function: A Historical Perspective. J Neurosci, 40(5), 944–954. doi:10.1523/JNEUROSCI.0740-19.2019

Patrie, K. M. (2005). Identification and characterization of a novel tight junction-associated family of proteins that interacts with a WW domain of MAGI-1. Biochim Biophys Acta, 1745(1), 131–144. doi:10.1016/j.bbamcr.2005.05.011

Piccoli, G., Verpelli, C., Tonna, N., Romorini, S., Alessio, M., Nairn, A. C.,…Sala, C. (2007). Proteomic analysis of activity-dependent synaptic plasticity in hippocampal neurons. J Proteome Res, 6(8), 3203–3215. doi:10.1021/pr0701308

Redmond, L., Kashani, A. H., & Ghosh, A. (2002). Calcium regulation of dendritic growth via CaM kinase IV and CREB-mediated transcription. Neuron, 34(6), 999–1010. doi:10.1016/s0896-6273(02)00737-7

Richier, L., Williton, K., Clattenburg, L., Colwill, K., O’Brien, M., Tsang, C.,…Fawcett, J. P. (2010). NOS1AP associates with Scribble and regulates dendritic spine development. J Neurosci, 30(13), 4796–4805. doi:10.1523/JNEUROSCI.3726-09.2010

Rojek, K. O., Krzemien, J., Dolezyczek, H., Boguszewski, P. M., Kaczmarek, L., Konopka, W.,…Proszynski, T. J. (2019). Amot and Yap1 regulate neuronal dendritic tree complexity and locomotor coordination in mice. PLoS Biol, 17(5), e3000253. doi:10.1371/journal.pbio.3000253

Schanzenbacher, C. T., Sambandan, S., Langer, J. D., & Schuman, E. M. (2016). Nascent Proteome Remodeling following Homeostatic Scaling at Hippocampal Synapses. Neuron, 92(2), 358–371. doi:10.1016/j.neuron.2016.09.058

Scudder, S. L., Goo, M. S., Cartier, A. E., Molteni, A., Schwarz, L. A., Wright, R., & Patrick, G. N. (2014). Synaptic strength is bidirectionally controlled by opposing activity-dependent regulation of Nedd4-1 and USP8. J Neurosci, 34(50), 16637–16649. doi:10.1523/JNEUROSCI.2452-14.2014

Sun, Q., & Turrigiano, G. G. (2011). PSD-95 and PSD-93 play critical but distinct roles in synaptic scaling up and down. J Neurosci, 31(18), 6800–6808. doi:10.1523/JNEUROSCI.5616-10.2011

Swann, J. W., Le, J. T., Lam, T. T., Owens, J., & Mayer, A. T. (2007). The impact of chronic network hyperexcitability on developing glutamatergic synapses. Eur J Neurosci, 26(4), 975–991. doi:10.1111/j.1460-9568.2007.05739.x

Szebenyi, G., Bollati, F., Bisbal, M., Sheridan, S., Faas, L., Wray, R.,…Brady, S. T. (2005). Activity-driven dendritic remodeling requires microtubule-associated protein 1A. Curr Biol, 15(20), 1820–1826. doi:10.1016/j.cub.2005.08.069

Thanh Nguyen, H., Andrejeva, D., Gupta, R., Choudhary, C., Hong, X., Eichhorn, P. J.,…Cohen, S. M. (2016). Deubiquitylating enzyme USP9x regulates hippo pathway activity by controlling angiomotin protein turnover. Cell Discov, 2, 16001. doi:10.1038/celldisc.2016.1

Thines, L., Gorisse, L., Li, Z., Sayedyahossein, S., & Sacks, D. B. (2022). Calmodulin activates the Hippo signaling pathway by promoting LATS1 kinase-mediated inhibitory phosphorylation of the transcriptional coactivator YAP. J Biol Chem, 298(5), 101839. doi:10.1016/j.jbc.2022.101839

Todi, S. V., & Paulson, H. L. (2011). Balancing act: deubiquitinating enzymes in the nervous system. Trends Neurosci, 34(7), 370–382. doi:10.1016/j.tins.2011.05.004

Turrigiano, G. G., Leslie, K. R., Desai, N. S., Rutherford, L. C., & Nelson, S. B. (1998). Activity-dependent scaling of quantal amplitude in neocortical neurons. Nature, 391(6670), 892–896. doi:10.1038/36103

Vallejo, D., Codocedo, J. F., & Inestrosa, N. C. (2017). Posttranslational Modifications Regulate the Postsynaptic Localization of PSD-95. Mol Neurobiol, 54(3), 1759–1776. doi:10.1007/s12035-016-9745-1

Vucic, D., Dixit, V. M., & Wertz, I. E. (2011). Ubiquitylation in apoptosis: a post-translational modification at the edge of life and death. Nat Rev Mol Cell Biol, 12(7), 439–452. doi:10.1038/nrm3143

Wang, C., An, J., Zhang, P., Xu, C., Gao, K., Wu, D.,…Yu, L. (2012). The Nedd4-like ubiquitin E3 ligases target angiomotin/p130 to ubiquitin-dependent degradation. Biochem J, 444(2), 279–289. doi:10.1042/BJ20111983

Wang, Y., & Yu, F. X. (2024). Angiomotin family proteins in the Hippo signaling pathway. Bioessays, 46(8), e2400076. doi:10.1002/bies.202400076

Wells, C. D., Fawcett, J. P., Traweger, A., Yamanaka, Y., Goudreault, M., Elder, K.,…Pawson, T. (2006). A Rich1/Amot complex regulates the Cdc42 GTPase and apical-polarity proteins in epithelial cells. Cell, 125(3), 535–548. doi:10.1016/j.cell.2006.02.045

Widagdo, J., Guntupalli, S., Jang, S. E., & Anggono, V. (2017). Regulation of AMPA Receptor Trafficking by Protein Ubiquitination. Front Mol Neurosci, 10, 347. doi:10.3389/fnmol.2017.00347

Wigerius, M., Quinn, D., Diab, A., Clattenburg, L., Kolar, A., Qi, J.,…Fawcett, J. P. (2018). The polarity protein Angiomotin p130 controls dendritic spine maturation. J Cell Biol, 217(2), 715–730. doi:10.1083/jcb.201705184

Wigerius, M., Quinn, D., & Fawcett, J. P. (2020). Emerging roles for angiomotin in the nervous system. Sci Signal, 13(655). doi:10.1126/scisignal.abc0635

Yamada, T., Yang, Y., & Bonni, A. (2013). Spatial organization of ubiquitin ligase pathways orchestrates neuronal connectivity. Trends Neurosci, 36(4), 218–226. doi:10.1016/j.tins.2012.12.004

Yi, J. J., & Ehlers, M. D. (2005). Ubiquitin and protein turnover in synapse function. Neuron, 47(5), 629–632. doi:10.1016/j.neuron.2005.07.008

Yoon, S., Parnell, E., & Penzes, P. (2020). TGF-beta-Induced Phosphorylation of Usp9X Stabilizes Ankyrin-G and Regulates Dendritic Spine Development and Maintenance. Cell Rep, 31(8), 107685. doi:10.1016/j.celrep.2020.107685

